# (Pro)Renin Receptor Antagonism Attenuates High-Fat-Diet–Induced Hepatic Steatosis in Non-Alcoholic Fatty Liver Disease

**DOI:** 10.1101/2022.09.07.506819

**Authors:** Ariana Julia B. Gayban, Lucas Souza, Silvana G. Cooper, Erick Regalado, Robert Kleemann, Yumei Feng Earley

## Abstract

Non-alcoholic fatty liver disease (NAFLD) comprises a spectrum of liver damage directly related to diabetes, obesity, and metabolic syndrome. The (pro)renin receptor (PRR) has recently been demonstrated to play a role in glucose and lipid metabolism. Here, we hypothesized that inhibition of the PRR would prevent the development of diet-induced hepatic steatosis and fibrosis. To test our hypothesis, we fed wild-type mice on a C57Bl/6J background either a high-fat diet (HFD; 60% calories from fat) or normal fat diet (NFD; 10% calories from fat) with matching calories for 6 weeks. An 8-week methionine choline-deficient (MCD) diet was used to induce fibrosis in C57BL/6J mice. Two weeks following diet treatment, mice were implanted with a subcutaneous osmotic pump delivering either PRO20, a peptide PRR antagonist, or scrambled peptide (700 μg/kg/d) for 4 or 6 weeks. We found that a 6-week HFD significantly increased liver lipid accumulation, as detected by Oil Red O staining, and liver triglyceride content compared with NFD-fed mice. Importantly, PRO20 treatment significantly reduced hepatic lipid accumulation in HFD-fed mice without affecting body weight or glucose levels. Furthermore, PRR antagonism attenuated HFD-induced steatosis, particularly microvesicular steatosis. In the MCD diet model, the percentage of collagen area detected by Sirius Red staining was reduced in PRO20-treated compared with control mice. PRO20 treatment also significantly decreased levels of liver alanine aminotransferase (ALT), an indicator of liver damage, in MCD-fed mice compared with controls. Mechanistically, we found that PRR antagonism prevented HFD-induced increases in PPARγ and glycerol-3-phosphate acyltransferase 3 expression in the liver. Taken together, our findings establish the mechanism by which PRR regulates lipid metabolism in the liver and suggest the therapeutic potential of PRR antagonism for the treatment of liver steatosis and fibrosis development in NAFLD.

## Introduction

Non-alcoholic fatty liver disease (NAFLD), comprising a spectrum of pathological liver disease stages, including liver cell damage, is closely associated with diabetes, metabolic syndrome, and obesity. In light of the prevalence of obesity and alarming growth in type 2 diabetes frequency, this condition presents a rising health threat in Western countries, where its incidence ranges from 20% to 30% and continues to grow, especially within the pediatric population [1]. Prominent hallmarks of this condition include an initial abnormal accumulation of triglycerides in hepatocytes resulting from, among other things, decreased β-oxidation, increased *de novo* lipogenesis, and increased free fatty acid flux from adipose tissue [2] in the absence of excessive alcohol intake. The pathogenesis of this disease is marked by a gradual loss of the liver to metabolize fatty acids and carbohydrates, leading to an abnormal accumulation of lipids in lipid droplets within hepatocytes, otherwise known as fatty liver. The recurrent instigation of hepatic steatosis is believed to increase the susceptibility of the liver to more severe forms of liver damage, typically in the form of non-alcoholic steatohepatitis (NASH), characterized by steatosis in conjunction with inflammation, which may present with increasingly severe forms of hepatic fibrosis [3]. Progression of this disease is accompanied by irreversible liver damage and life-threatening complications, including hepatocellular carcinoma and, ultimately, liver failure. Current treatments for NAFLD are ineffective in preventing disease progression towards NASH and fibrosis, underscoring the need for the development of novel and potent treatment methods as well as the identification of valid drug targets.

One novel target worth investigating for the treatment of NAFLD is the (pro)renin receptor (PRR). The PRR, a single transmembrane receptor broadly expressed in the kidneys, heart, vascular smooth muscle, brain, adipose tissue, liver and eye [5–10], is a key component of the renin-angiotensin system (RAS) that is involved in regulating blood pressure through angiotensin II-dependent and angiotensin II-independent pathways [4]. Investigations of this receptor in various tissues have implicated prorenin-mediated PRR function in the development of features of metabolic syndrome, such as diabetes, obesity, and obesity-related hypertension [11–13]. In addition, several studies have shown that the PRR is an emerging player in the regulation of lipid metabolism [9, 14], suggesting that the PRR could be a potential novel target for the treatment of metabolic syndrome-related diseases, including non-alcoholic fatty liver. A role for the PRR in regulating plasma lipids through integration of glucose and lipid metabolism in the liver, as well as LDL metabolism through a low-density lipoprotein receptor (LDLR)- and sortilin-1 (SORT1)-mediated mechanism, has recently been demonstrated [15]. The PRR has also recently been shown to play a role in mediating the onset of fibrosis in the context of kidney pathology [16, 17]. However, the molecular mechanisms underlying the involvement of the PRR in fatty liver development, hepatic fibrogenesis, and hepatic triglyceride metabolism have yet to be elucidated.

In the current study, we used PRR antagonism, employing the 20-amino-acid peptide, PRO20, to investigate the role of the PRR in modulating liver lipid metabolism in the setting of NAFLD. PRO20, which consists of the first 20 amino acids of the (pro)renin prosegment [18], displays high specificity for both the human and mouse PRR and outcompetes (pro)renin for PRR binding sites. The use of PRO20 as a tool for studying the action of PRR in the pathogenesis of disease has been validated previously in mouse models [18–20]. In the current study, we demonstrate the action of PRR as a novel regulator of hepatic triglyceride metabolism and a potential therapeutic tool against in the development of fatty liver and NASH.

## Materials and Methods

### Animals

Male 16-week-old C57BL/6J mice were obtained from Jackson Laboratories (strain#: 000664). Mice were housed individually and fed either a high-fat diet (HFD; D12492 Research Diets Inc.) or normal-fat diet (NFD; D12450J Research Diets Inc.) containing 60% and 10% of kCal from fat, respectively, for 6 weeks. An 8-week methionine choline-deficient (MCD) diet (TD.90262, Envigo) regimen was used to induce a NASH phenotype in 16-week-old male C57BL/6J mice under the same single-housed conditions. Body weight and food intake were measured for all mice weekly. Following the diet modification, mice were sacrificed, and tissues were collected for molecular experiments. All animal procedures were approved by the Institutional Animal Care and Use Committee at the University of Nevada, Reno, and were performed in accordance with the National Institutes of Health Guidelines for the care and use of experimental animals.

### Subcutaneous Administration of PRO20 (or Scrambled Peptide) or Losartan (or Saline) Using Osmotic Minipumps

After 2 weeks of diet treatment (NFD or HFD), mice received subcutaneous osmotic minipump implants (Alzet Micro-Osmotic Pump Model 1004) to administer the PRO20 peptide (LPTRTATFERIPLKKMPSVREI) or scrambled peptide (control, LRTETPITMIPSAERVFRKKPL) at 700 μg/kg/d for the remaining 4 weeks of treatment. For the MCD diet, mice received a similar osmotic minipump implant (Alzet Micro-Osmotic Pump Model 2006) after 2 weeks of MCD treatment. These pumps administered the same dose of either PRO20 or scrambled peptide over the remaining 6 weeks of diet treatment. Mice were anesthetized by isoflurane inhalation and then subcutaneously implanted with osmotic minipumps infusing PRO20 or scrambled (control) peptide. A separate study using losartan, an angiotensin II type 1a receptor (AT1aR) blocker, was performed using the aforementioned 6-week HFD and NFD diet treatment regimen. Specifically, after 2 weeks of diet treatment, mice received subcutaneous osmotic minipump implants that delivered either losartan (10 mg/kg/d) or 0.9% saline (control) for the subsequent 4 weeks. All animals were housed singly in standard forced-air shoebox cages and were maintained on their respective diets until the end of the treatment period.

### Fasting Blood Glucose Measurement and Glucose Tolerance Tests

Fasting blood glucose (FBG) levels were measured in all mice at baseline and after 6 weeks of either a NFD or HFD regimen. Mice were fasted for 16 hours (6 PM to 10 AM) in clean cages before glucose measurements. Blood was collected from each mouse by creating a 1-mm cut at the tip of the tail, and subsequent FBG values were measured using a Bayer 7393A Contour blood glucose meter. Glucose was measured in duplicate for each mouse, and the mean of the two values was taken as the final measurement.

Glucose tolerance tests (GTTs) were performed before the beginning of treatment and after 6 weeks of diet modification. Mice were fasted for 16 hours (6 PM to 10 AM) in clean cages prior to beginning the GTT. After fasting, baseline blood glucose was measured and a solution of 10% glucose in 0.9% sterile saline (1 g/kg body weight) was injected intraperitoneally to elevate blood glucose levels. For GTTs, blood glucose was measured and recorded for each mouse 15, 30, 60, 90, and 120 minutes after injection of glucose using a Bayer blood glucose meter, as described above.

### Oil Red O Staining

At the end of the 6-week study, all mice were sacrificed and liver tissue was processed to obtain frozen liver cross-sections as well as paraffin-embedded cross-sections. The left lobe of the liver was kept in 4% paraformaldehyde for 24 hours, then in 30% sucrose for the following 24 hours, and subsequently frozen in tissue freezing medium (Tissue Freezing Medium, GeneralData TFM-Y). Serial frozen liver sections were cut at 10 μm thickness using a cryostat (Leica Biosystems). Liver sections were affixed to glass slides by air drying and then fixed in formalin for 5 minutes. After a 1-minute wash in tap water, slides were rinsed in 60% isopropanol and stained with Oil Red O solution (Abcam, ab150678) for 30 minutes. After staining, slides were rinsed in 60% isopropanol and counterstained with Mayer Hematoxylin (1 g/L; Sigma Aldrich, MHS32) for 10 seconds, and then mounted. Images were captured using a light microscope (BZX-710; Keyence), and the percent area of Oil Red O staining was measured using ImageJ/FIJI.

### Hematoxylin & Eosin Staining

Hematoxylin & Eosin (H&E) staining was performed on paraffin-embedded median lobes of the liver. Paraffin-embedded liver tissues were cut at a thickness of 10 μm using a microtome (AccuCut; Tissue-Tek). Slides were rehydrated using decreasing graded series of ethanol concentrations (100%, 95%, 80%, 70%), then stained with Harris Hematoxylin (Lerner, 1931382) and Eosin Y (Sigma, H911032). Slides were dehydrated with an increasing graded series of ethanol concentrations (50%, 70%, 80%, 95%, 100%) and mounted. Images were acquired under a light microscope (Keyence BZX-710) and used for further analysis of liver grade, microvesicular and macrovesicular steatosis, and cardiomyocyte size.

### Assignment of Histological NAFLD Score and Measurement of Cardiomyocyte Diameter

A previously developed rodent NAFLD grading system [21] was used to score the extent of overall NAFLD development in each treatment group and the severity of hepatocellular vesicular steatosis (i.e., both microvesicular and macrovesicular steatosis). Microvesicular steatosis was defined as the presence of hepatocellular lipid vacuoles that did not displace the nucleus to the side, whereas macrovesicular steatosis featured the presence of large lipid vacuoles that displaced the nucleus. The severity of both types of vesicular steatosis was based on the total area of the slide affected, based on analysis of images taken at 20x magnification. Scores were assigned from least to most severe based on the percentage of area affected, as follows: 0, <5%; 1,5%–33%; 2, 34%–66%; and 3, >66%. The sum of the scores derived from both microvesicular and macrovesicular area percentages was considered the total steatosis grade.

### Trichrome Staining

Paraffin-embedded sections of the median lobe of the liver, cut at 10 μm, were rehydrated with descending concentrations of ethanol and subsequently washed in deionized water. Slides were then incubated in Bouin’s solution (Sigma) at 56°C for 15 minutes and then in Weigert’s Hematoxylin for 5 minutes, with 5-minute washes in deionized water between and after the two staining procedures. Slides were incubated sequentially in a Biebrich Scarlet-Acid Fuschin solution, phosphotungisic/phosphomolybdic acid solution, and aniline blue solution for 5 minutes each. Thereafter, slides were placed in 1% acetic acid for 2 minutes and then dehydrated in an ascending graded ethanol series (50%, 70%, 80%, 95%, 100%). Samples were cleared with xylene and mounted. Trichrome staining was observed and imaged under a light microscope (Keyence, BZX-710), and the area of blue collagen staining, as a percentage, was measured using ImageJ/FIJI.

### Picrosirius Red Staining

Paraffin-embedded sections of the median lobe of the liver were cut at a thickness of 10 μm and then stained with Picrosirius Red, which supplements Masson’s Trichrome staining in assessing hepatic fibrosis. Slides were incubated in xylene and then rehydrated by soaking in a descending graded series of ethanol concentrations (100%, 95%, 80%, 70%). Samples were then washed in phosphate-buffered saline (PBS) for 3 minutes, followed by an 8-minute incubation in Weigert’s hematoxylin and 10-minute wash in tap water. The tissue was then stained in Sirius Red solution for 1 hour, after which slides were immersed in two changes of acidified water for 2 minutes each. Slides were then dehydrated in 3 changes of 100% ethanol for 2 minutes each, cleared in xylene for 5 minutes, and then mounted. Images were captured using a light microscope (Keyence BZX-710), and the area of red collagen staining, expressed as a percentage, was measured using ImageJ/FIJI.

### Liver Alanine and Aspartate Aminotransferase Activity Assay

Liver injury in MCD-treated mice was assessed biochemically by quantifying plasma alanine (AST) and aspartate aminotransferase (ALT) activity. Colorimetric assays for AST (Sigma-Aldrich, MAK055) and ALT (Sigma-Aldrich, MAK052) activity levels were performed according to the manufacturer’s protocols. Briefly, plasma samples were added to microplate wells containing prepared AST or ALT reaction mixes. Optical density (OD) of the samples was measured at a wavelength of 450 nm for AST and 570 nm for ALT using a microplate reader (FlexStation 3). Plates were repeatedly incubated at 37°C for 5 minutes, and OD measurements were taken following each incubation until the value of the most active sample eclipsed the value of the highest standard. AST and ALT activity was calculated in milliunits/mL using a standard curve generated for each assay, assay reaction times, and the difference between the final and initial OD measurement.

### Quantitative Reverse Transcription-Polymerase Chain Reaction (qRT-PCR)

Total mRNA was extracted from frozen liver samples using the Trizol solubilization and extraction method according to the manufacturer’s protocol (ThermoFisher, 15596018). The isolated RNA was resuspended in 50 μL of ultra-pure water, and the quality and yield of the total RNA was determined using a Nanodrop spectrophotometer.

RNA contaminants were eliminated using a DNAse I treatment kit (ThermoFisher BP81071) and cDNA was produced from total RNA by reverse transcription using a High-Capacity cDNA reverse transcription kit (ThermoFisher, 4368814). Real-time qPCR was performed on a Quantstudio 3 System in 20-μL reactions using SYBR Green PCR Master Mix (ThermoFisher). mRNA levels of the following targets were measured and reported as fold-change in mRNA expression, determined using the ΔΔCT method: peroxisome proliferator activated receptor (PPAR)-α, -β, and -γ; β-actin; carbohydrate-responsive element-binding protein (CHREBP); sterol regulatory element-binding protein 1c (SREBP1c); acetyl-CoA carboxylase (ACC); fatty acid synthase (FAS); ATP-citrate lyase (ATPCL); and glycerol-3-phosphate acyltransferase 3 (GPAT3). The primer sequences used for each gene are listed in Table 1.

**Table 1.**
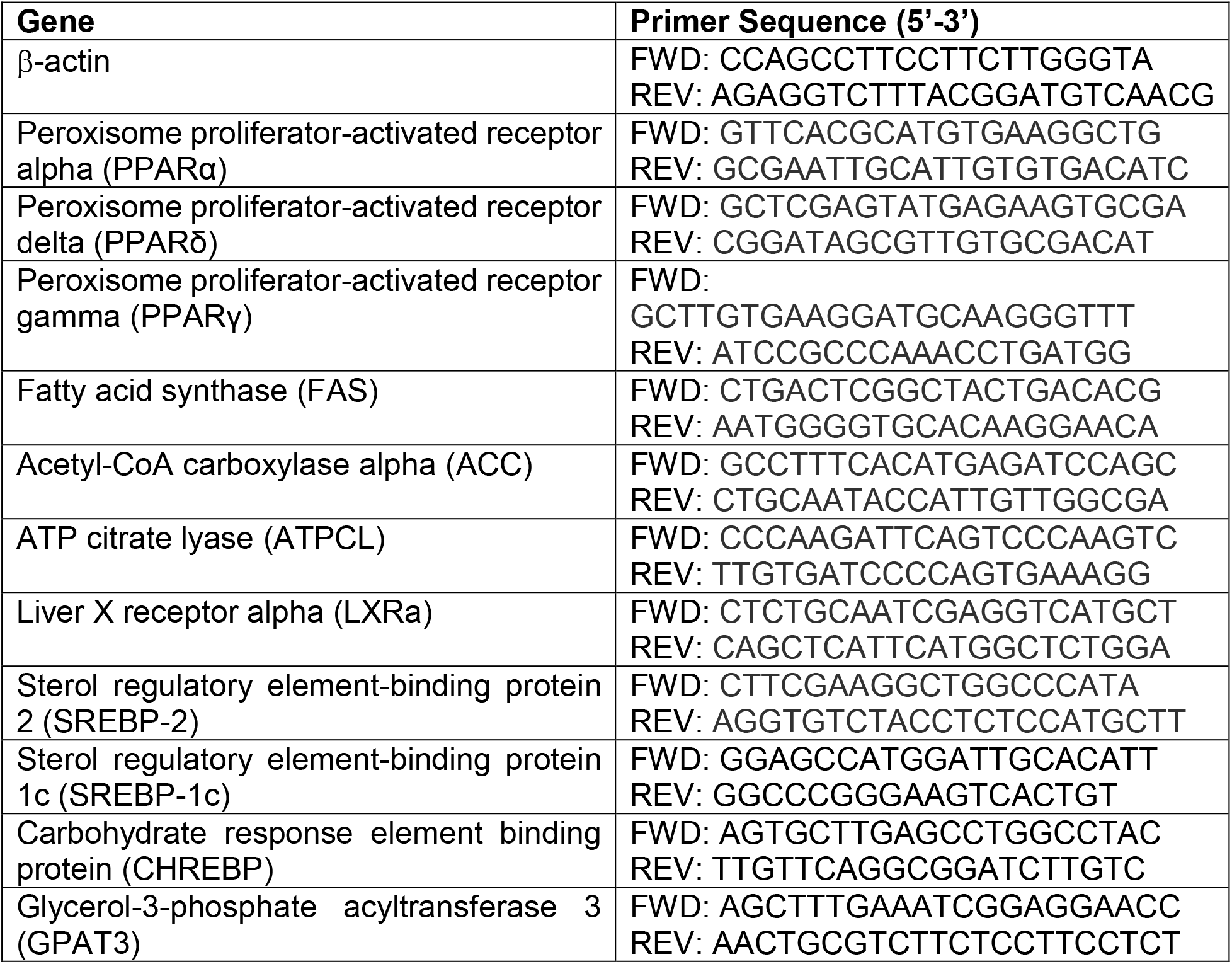
Sequences of primers used in the qPCR analysis of PPARs and triglyceride synthesis components.

### Western Blot Analysis

Protein abundance of β-actin and GPAT3 was investigated by Western blot analysis. Liver total protein was extracted using RIPA lysis buffer (ThermoFisher, catalog# 89901) and protein concentrations were determined using BCA assays (ThermoFisher, catalog# 23225). Protein-containing samples were prepared for sodium dodecyl sulfatepolyacrylamide gel electrophoresis (SDS-PAGE) by denaturation in a solution containing 2% SDS and 100 mM DTT at 95°C for 5 minutes followed by centrifugation at 13,000 × g for 10 minutes. Equal amounts of protein (10–30 μg) were resolved on 4–12% Tris-Glycine gels (Invitrogen, NW04125BOX). After gel electrophoresis, proteins were transferred onto a nitrocellulose membrane using a traditional wet transfer method for 70 minutes at 4°C. Blots were then blocked by incubating in Tris-buffered saline containing 2.5% BSA for 1 hour at room temperature. After blocking, blots were incubated with rabbit anti-mouse β-actin (1:1000 dilution; Cell Signaling Technology 8457S) or rabbit antimouse GPAT3 (1:500 dilution; ThermoFisher, catalog# 20603-1-AP) primary antibody. Following incubation with primary antibody, blots were transferred to a solution containing anti-rabbit IgG HRP-linked secondary antibody (1:1000 dilution; Cell Signaling Technology 7074S) and incubated for 1 hour at room temperature. Immunoreactive proteins were detected using SuperSignal West Dura Extended Duration Substrate (ThermoFisher, catalog# 34076) in conjunction with the BioRad ChemiDoc MP Imaging System. Band intensity was quantified using ImageJ/FIJI software.

## 3. Results

### PRR Antagonism Attenuates HFD-Induced Lipid Accumulation in the Liver

To determine whether the PRR regulates hepatic lipid metabolism and plays a role in the development of hepatic steatosis, we used the PRR antagonist, PRO20, to block PRR activation in a HFD-induced mouse model of NAFLD. PRR antagonism reduced fat accumulation in the liver, as detected by Oil Red O staining (Figure 1A). Quantitative analyses showed that, after a period of 6 weeks, HFD induced a significant accumulation of fat (16.6% ± 1.0%) compared with NFD controls (3.3% ± 0.4%, *P* < 0.00001). This effect of HFD was attenuated by administration of PRO20, which significantly decreased hepatic fat accumulation (7.7% ± 1.4%, *P* < 0.00001) compared with HFD controls (Figure 1B), albeit without completely normalizing fat content compared with mice that received NFD and scrambled peptide treatment. PRO20 had no effect on lipid accumulation in NFD-fed mice.

**Figure 1.**
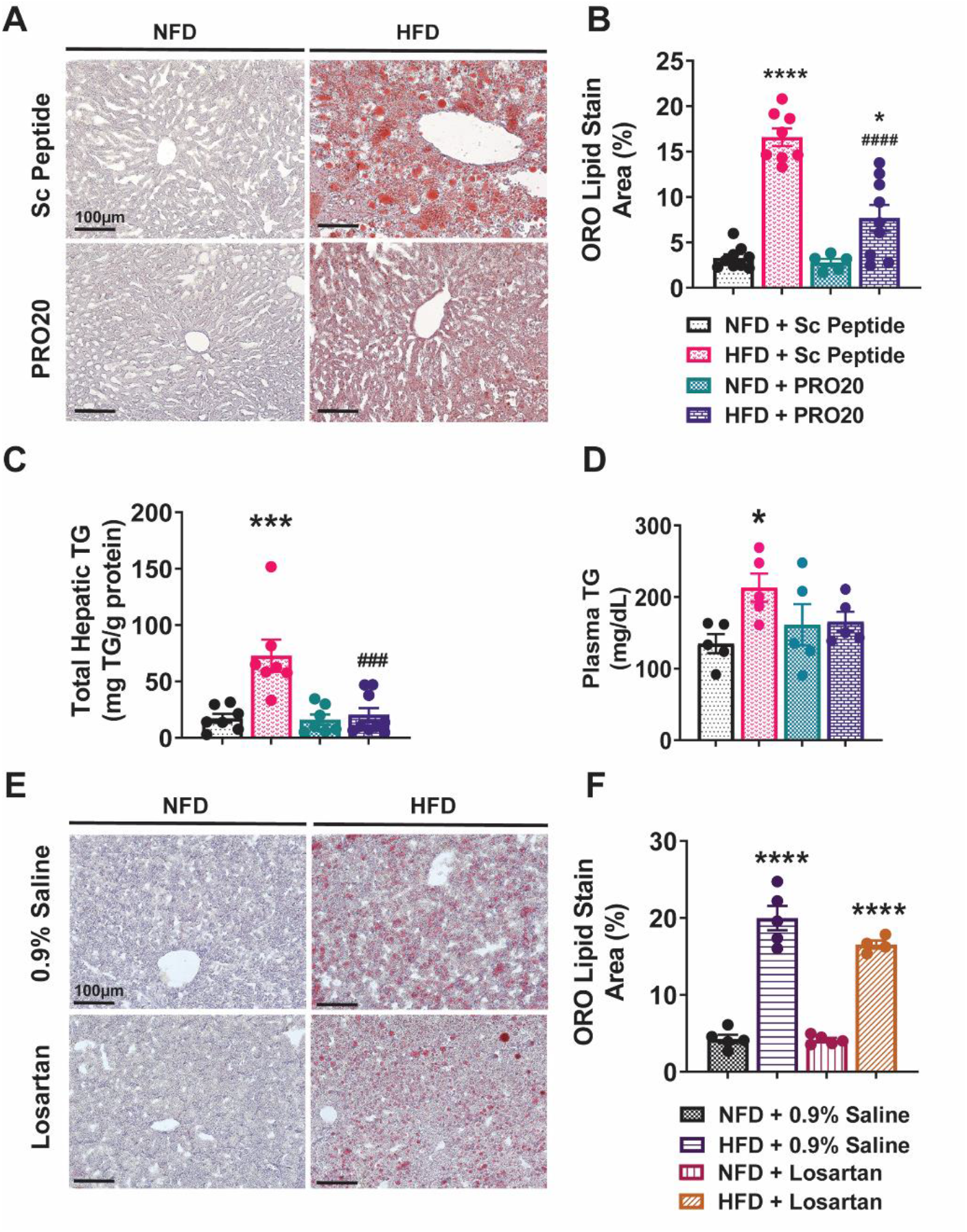
The PRR antagonist, PRO20, but not losartan, reduces hepatic steatosis under HFD conditions. **(A)** Representative images of hepatic lipid accumulation, visualized by Oil Red O (ORO) staining in mice from PRO20 and control groups fed either a HFD or NFD. **(B)** Quantification of ORO staining, presented as the percentage area of red staining relative to total image area. **(C)** Biochemical assay of total hepatic triglycerides (TG) normalized to total liver protein levels. **(D)** Biochemical assessment of plasma TG concentrations in mice. **(E)** Representative images of hepatic lipid accumulation, determined by ORO staining in mice from losartan and control groups under HFD or NFD conditions. **(F)** Quantification of ORO staining from the losartan study presented as the area of red staining relative to the total image area, expressed as a percentage. Data are presented as means ± SEM (\**P* < 0.05, \*\*\**P* < 0.001, \*\*\*\**P* < 0.0001 vs. NFD + Sc Peptide or NFD + 0.9% Saline; ###*P* < 0.001, ####*P* < 0.0001 vs. HFD + Sc Peptide; one-way ANOVA).

To further confirm the lipid-accumulation status in the liver, we measured total hepatic triglyceride levels and normalized them to total protein present in each sample. As shown in Figure 1C, 6 weeks of HFD feeding induced a significant increase in total hepatic triglycerides (72.9 ± 14.2 mg/g protein) in mice that received the scrambled peptide. Notably, this effect was blunted by administration of PRO20, which reduced the concentration of hepatic triglycerides (20.5 ± 6.0 mg/g protein) compared with HFD controls. These data confirm Oil Red O results showing that HFD treatment induces lipid accumulation and that PRR antagonism alleviates this accumulation. The increase in hepatic triglycerides in HFD-fed mice was accompanied by a significant increase in plasma triglyceride levels (Figure 1D). Specifically, circulating triglycerides increased from 134.9 ± 13.3 mg/dL in NFD-fed mice to 213.1 ± 19.8 mg/dL in mice fed HFD for 6 weeks (*P* = 0.0137). Although plasma triglyceride levels trended lower in HFD-fed mice treated with PRO20 (165.6 ± 13.94 mg/dL), this decrease did not achieve statistical significance (*P* = 0.1123).

### Subcutaneous Infusion of Losartan Does Not Reduce Hepatic Lipid Accumulation in HFD-fed Mice

Activation of the PRR mediates formation of angiotensin (Ang) peptides, as well as stimulation of Ang II-independent signaling pathways [4, 22]. To investigate the extent to which Ang II/AT1R signaling activation impacts the development of non-alcoholic fatty liver, we administered the AT1R antagonist, losartan, using the same protocol as above for PRO20. Representative images of Oil Red O-stained tissues from NFD- or HFD-fed mice treated with losartan or 0.9% saline are shown in Figure 1E. As expected, 6 weeks of HFD induced an increase in liver lipid accumulation (20.0% ± 1.6%) relative to NFD-fed mice infused with saline (4.3% ± 0.6%, *P* < 0.0001). Hepatic lipid accumulation tended to be lower with subcutaneous losartan infusion for 4 weeks (16.5% ± 0.5%), but the Oil Red O-positive area was not significantly reduced compared with controls (20.0% ± 1.6%, *P* = 0.09). Thus, at the end of the 6-week HFD regimen (Figure 1F), HFD-fed mice that received losartan treatment still displayed an increase in hepatic lipid content compared with NFD-fed control mice (4.3% ± 0.6%, *P* < 0.0001). These data suggest that losartan does not attenuate HFD-induced lipid accumulation in the liver in our experimental time frame.

### PRO20 Attenuates HFD-Induced Hepatic Steatosis

To examine structural changes in the liver following HFD, we performed H&E staining (Figure 2A), applying a histological scoring system commonly used to gauge the severity of NAFLD to compare the extent of histological changes among groups. Using a 0-3 scale developed for liver grading in rodents described by Liang et al. [21], we found that HFD-fed mice scored consistently higher than their NFD-fed counterparts in overall NAFLD severity (Figure 2B). In contrast, the liver grading scores of HFD-fed mice administered PRO20 were significantly lower than those of HFD controls and on par with those of NFD-fed mice.

**Figure 2.**
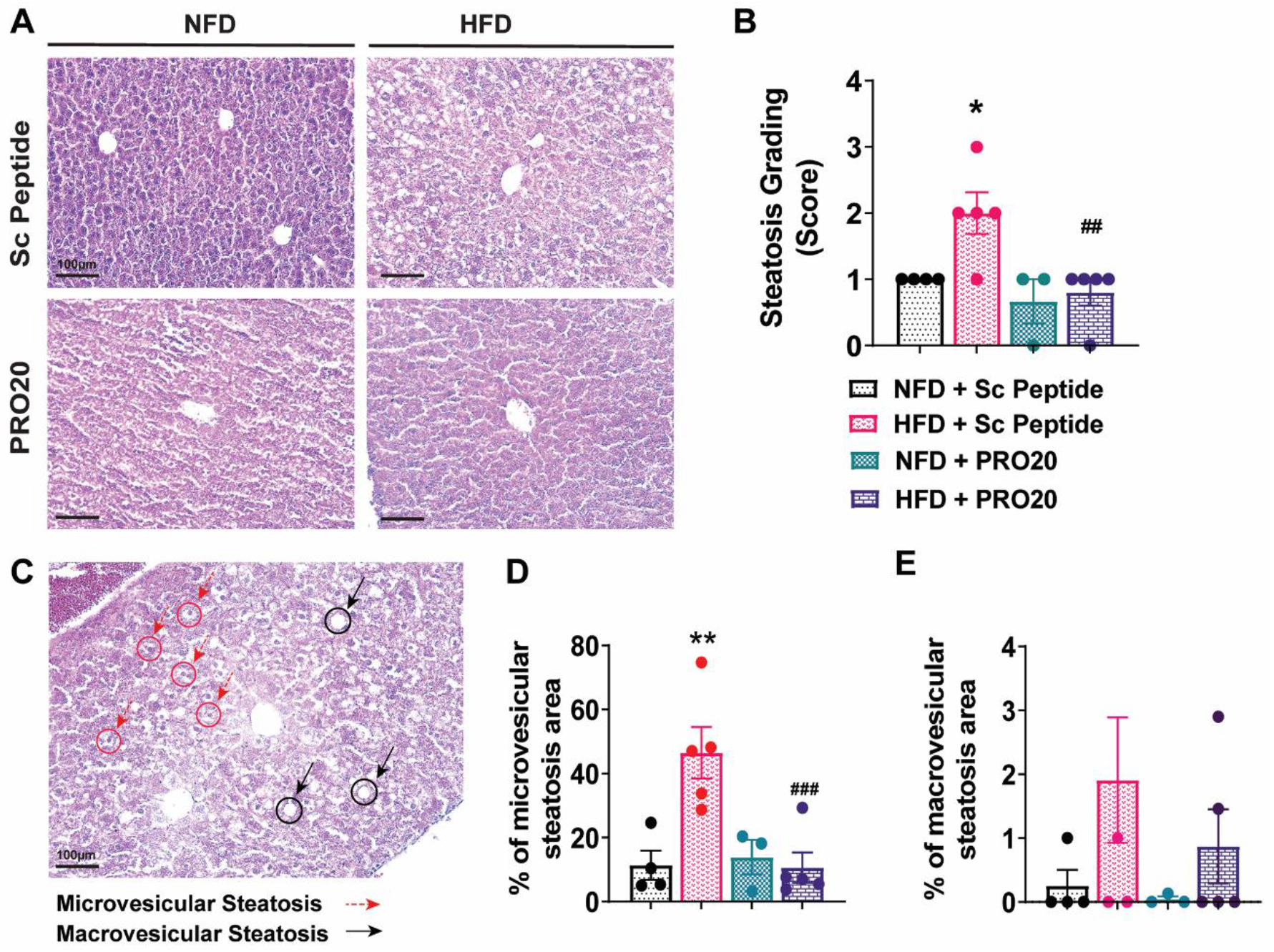
The PRR antagonist PRO20 decreases hepatic steatosis severity and microvesicular steatosis abundance. **(A)** Representative images of H&E-stained hepatic tissue, showing differences in tissue structure among treatment groups. **(B)** Distribution of liver steatosis grade in mice. Scores (0-3) were assigned based on the presence of microvesicular and macrovesicular steatosis, with 0 being the least severe and 3 corresponding to maximum steatosis severity. **(C)** Examples of microvesicular steatosis (dashed red arrows) and macrovesicular steatosis (solid black arrows) in mouse liver tissue. **(D, E)** Quantification of microvesicular (D) and macrovesicular (E) steatosis area, reported as microvesicular/macrovesicular lipid area relative to total image area, expressed as a percentage (n = 3–5 mice/group). Data are presented as means ± SEM (\**P* < 0.05, \*\**P* < 0.01 vs. NFD + Sc Peptide; ##*P* < 0.01, ###*P* < 0.001 vs. HFD + Sc Peptide; one-way ANOVA).

Next, we evaluated characteristic histological features of fatty liver disease, focusing on the development of microvesicular steatosis and macrovesicular steatosis [21]. As shown in Figure 2C, HFD induced an increase in microvesicular (red circles) and macrovesicular (black circles) steatosis as a result of increased hepatic lipid, with the area occupied by microvesicular steatosis reaching 46.5% ± 8.0% in HFD-fed mice compared with 11.4% ± 4.6% in NFD controls (*P* = 0.0013). Notably, we found a significant reduction in the severity of microvesicular steatosis in HFD-fed mice following treatment with the PRR antagonist PRO20, which reduced the total steatosis area to 10.6% ± 4.7% (*P* = 0.0007) (Figure 2D). Six weeks of HFD did not significantly increase macrovesicular steatosis compared with NFD-fed mice, and there were no differences in macrovesicular steatosis among treatment groups (Figure 2E).

### Development of Hepatic Fibrosis In Mice Fed a 6-Week HFD

Dysregulation of lipid metabolism in the liver has been shown to initially manifest as formation of steatosis (NAFLD), whereas advanced stages of liver damage are characterized by the additional development of hepatic inflammation (NASH), which may present with and without fibrosis [1]. Accordingly, we investigated whether a 6-week HFD induced fibrosis concurrent with abnormal lipid accumulation, and whether PRR antagonism could rescue this phenotype. Quantification of collagen in the liver with Masson’s trichrome stain showed that mice fed a 6-week HFD exhibited an increase in collagen development around the hepatic portal vein (4.1% ± 0.4%) compared with NFD controls (2.7% ± 0.4%) (Figure 3A and 3B). Notably, treatment of HFD-fed mice with PRO20 reduced the collagen area (1.7% ± 0.3%, P = 0.0007) compared with that in control HFD-mice administered a scrambled peptide. These results were further validated by Picrosirius Red staining, which employs an acidic dye that enhances the birefringence of collagen, allowing for more detailed visualization of polarized light compared with Masson’s Trichrome stain, which utilizes an aniline blue dye that binds to basic sites in collagen molecules [23]. Using this approach, we demonstrated a significant increase in hepatic collagen development in mice fed a 6-week HFD (5.3% ± 0.6%) compared with NFD controls (3.5% ± 0.3%, *P* = 0.0188) (Figure 3C and 3D), supporting our original findings. HFD-fed mice administered PRO20 also displayed a trend toward a reduction in collagen quantity, but this decrease was not significant. To complement histological analyses, we quantified the total hepatic collagen content in liver homogenates using a biochemical assay (Figure 3E). This biochemical assessment showed no significant difference in collagen content among treatment groups. In any case, the overall collagen quantity among groups, measured either histologically or biochemically, remained minimal, indicating that steatosis and not fibrosis is the predominant form of liver pathology present in this 6-week HFD model.

**Figure 3.**
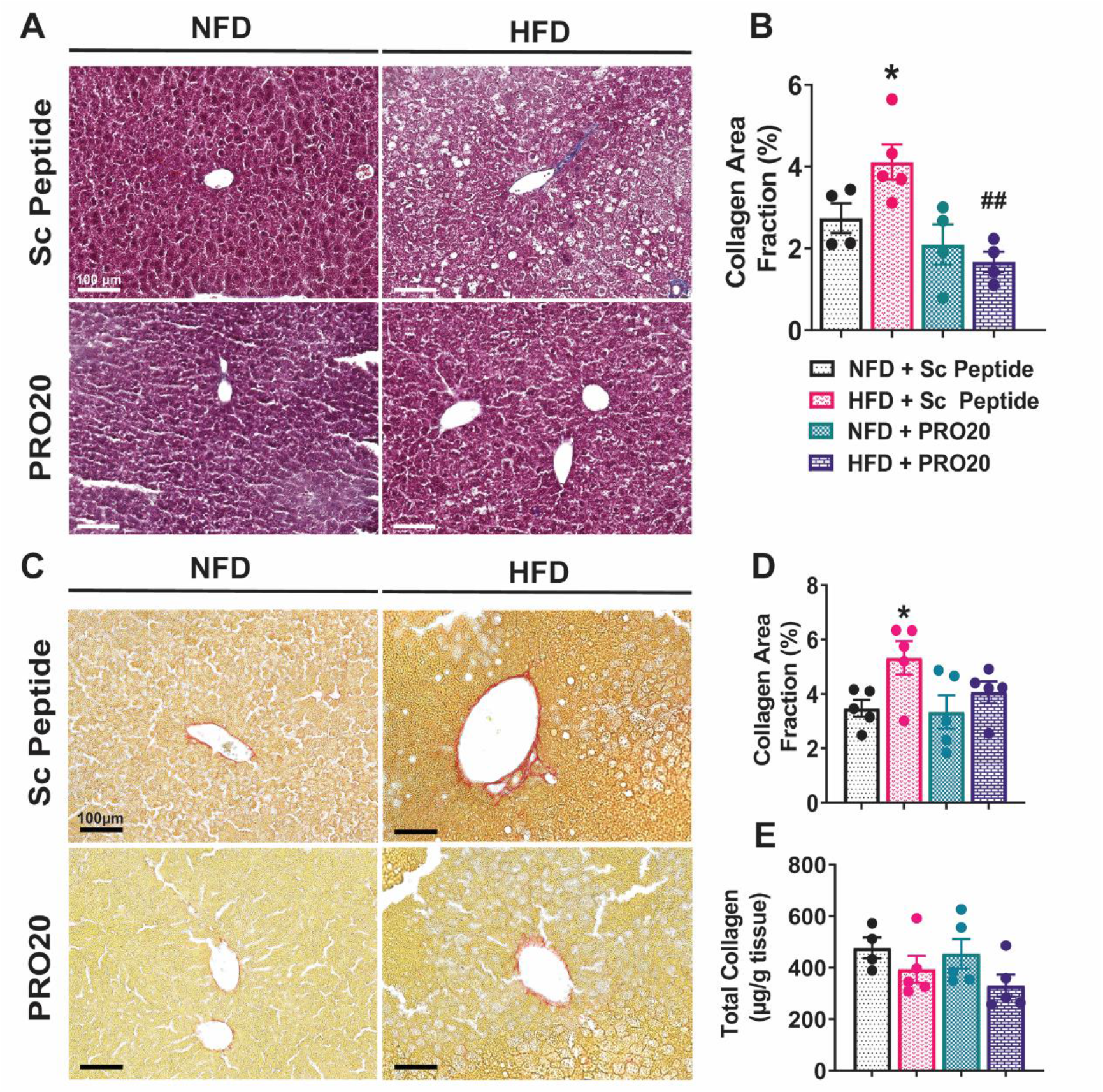
Six weeks of HFD promotes an increase in hepatic collagen deposition. **(A)** Representative images of hepatic collagen deposition, visualized by Masson’s trichrome staining. **(B)** Quantification of Masson’s trichrome collagen staining following 6 weeks of HFD or NFD treatment and either PRO20 or Sc Peptide administration. Data are presented as blue collagen staining relative to total image area, expressed as a percentage. **(C)** Representative images of hepatic collagen deposition visualized by Picrosirius Red stain. **(D)** Quantification of Picrosirius Red collagen staining. Data are presented as red collagen staining relative to total image area, expressed as a percentage. **(E)** Hydroxyproline-based biochemical analysis of total hepatic collagen normalized to tissue quantity. Data are presented as means ± SEM (\**P* < 0.05 vs. NFD + Sc Peptide; ##*P* < 0.01 vs. HFD + Sc Peptide; one-way ANOVA).

### PRO20 Decreases Hepatic Fibrosis Development and Liver Injury in MCD Diet-Fed Mice

To investigate the role of PRR in a more pronounced model of hepatic fibrosis, we employed an MCD diet model and administered PRO20 and scrambled peptides in tandem, as depicted in the protocol shown in Figure 4A. Body weight and food intake parameters were monitored weekly following the start of the MCD diet in all mice. No significant differences in body weight or food intake were observed between PRO20 and scrambled peptide treatment groups fed an MCD diet (Figure 4B); however, mice in both groups experienced a large drop in body weight, which is characteristic of this diet model [24]. Following an 8-week MCD diet and infusion of PRO20 or scrambled peptide for 6 weeks, livers were assessed for collagen deposition indicative of fibrosis development using Picrosirius Red staining and a biochemical collagen assay. Representative images (Figure 4C) and quantification of Picrosirius Red-stained area (Figure 4D) showed that PRO20 significantly reduced collagen deposition following MCD treatment, reducing the total area to 6.0% ± 0.6% compared with 8.9% ± 0.2% for scrambled peptide (*P* < 0.0001), indicating that the PRR may regulate the onset of liver fibrosis in MCD diet-induced NASH. Plasma ALT and AST levels, markers of liver injury, were also assessed in both PRO20- and scrambled peptide-treated mice (Figure 4E and 4F). PRO20 treatment significantly decreased circulating ALT activity (422.3 ± 65.2 U/L) compared with scrambled peptide controls (683.1 ± 83.41 U/L, *P* = 0.0249), but had no effect on AST activity.

**Figure 4.**
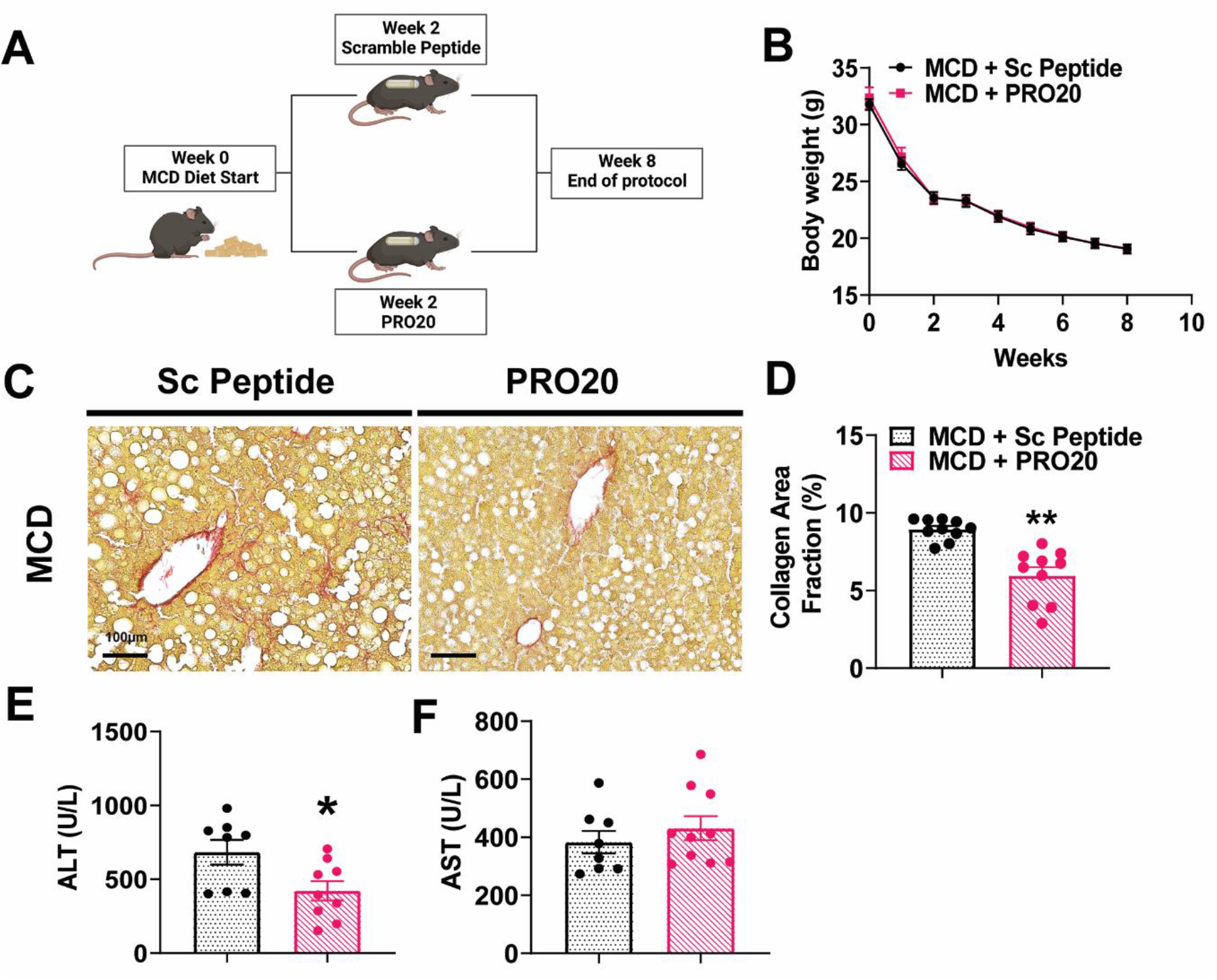
PRO20 treatment reduces hepatic fibrosis development and liver damage following 8 weeks of MCD Diet. **(A)** Schematic of MCD diet study protocol. **(B)** Summary data showing weekly body weight measurements in mice under MCD diet conditions. **(C)** Representative images of hepatic collagen deposition, visualized by Picrosirius Red staining. **(D)** Quantification of Picrosirius Red collagen stain. Data are presented as red collagen staining relative to total image area, expressed as a percentage. **(E, F)** Biochemical assessment of plasma ALT (E) and AST (F) levels. Data are presented as means ± SEM (\**P* < 0.05, \*\**P* < 0.01 vs. MCD + Sc Peptide; Unpaired t-test).

### PRR Antagonism Attenuates HFD-Induced Elevation of PPARγ and GPAT3 in the Liver

Because HFD-fed, PRO20-treated mice displayed significantly reduced hepatic triglyceride levels, we investigated whether genes involved in *de novo* lipogenesis in the liver were affected by the PRR antagonist PRO20. A 6-week HFD regimen, with or without PRO20, had no effect on mRNA expression levels of transcription factors involved in lipogenesis, including liver X receptor alpha (LXRα), carbohydrate response element binding protein (ChREBP), and sterol regulatory element-binding protein 1 (SREBP1c) (Figure 5A–C). As part of our investigation of changes in lipid homeostasis, we also assessed hepatic mRNA expression of members of the PPAR family of transcriptional regulators. The three known PPAR isoforms—PPARα, PPARδ, and PPARγ—have been shown to play a significant role in NAFLD pathology and liver physiology. In particular, activation of PPARα has been shown to promote fatty acid oxidation and transport, whereas the action of PPARγ leads to increased hepatic lipogenesis, triglyceride storage, and adipogenesis [25, 26]. PPARδ, though most abundant in muscle tissue, functions through interactions with the other PPAR isoforms to inhibit hepatic lipogenesis and insulin resistance [27, 28]. After a 6-week HFD, hepatic PPARγ was significantly upregulated (2.6 ± 0.6) compared with that observed in NFD controls (1.13 ± 0.19, *P* = 0.0037) (Figure 5D), an effect that was attenuated by administration of PRO20. Hepatic PPARα mRNA expression also trended higher in HFD-fed mice; although this increase did not reach statistical significance, it was significantly attenuated by PRO20 treatment (1.4 ± 0.1) compared to treatment with scrambled peptide (0.8 ± 0.1, P = 0.0040) (Figure 5E). PRO20 treatment also showed a tendency to reduce PPARδ mRNA levels in HFD-fed mice (1.0 ± 0.1) compared to HFD-fed mice treated with scrambled peptide (0.74 ± 0.07, P = 0.0319), although neither this difference nor the difference in PPARδ expression between HFD- and NFD-fed conditions were significant (Figure 5F).

**Figure 5.**
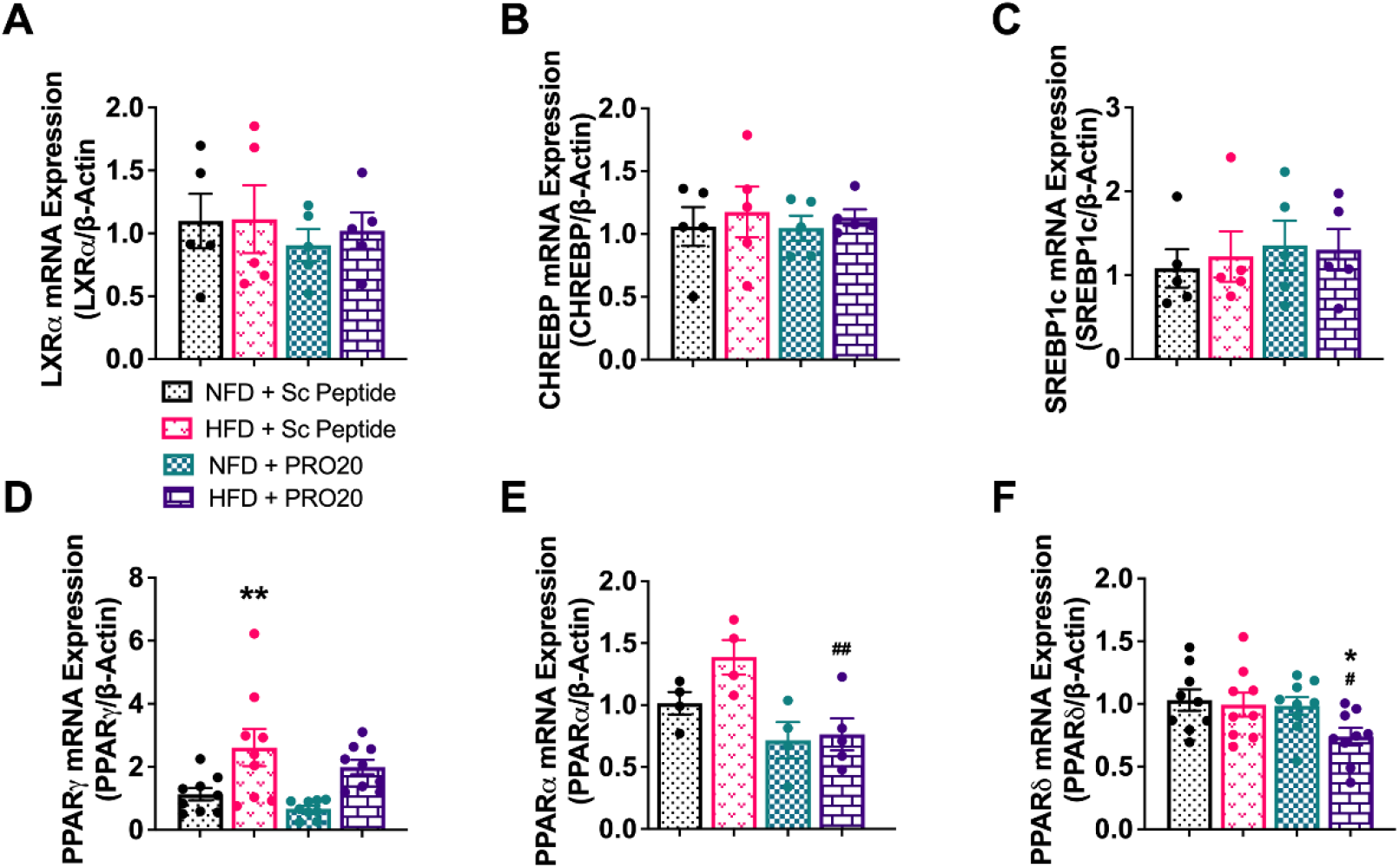
Six weeks of HFD increases hepatic mRNA expression of lipogenic PPARγ. Hepatic mRNA levels of the transcription factors LXRα **(A)**, CHREBP **(B)**, SREBP1c **(C)**, PPARγ **(D)**, PPARα **(E)**, and PPARδ **(F)** following 6 weeks of HFD or NFD and either PRO20 or Sc Peptide treatment. mRNA expression was assessed by RT-qPCR and was normalized to β-actin. Data are presented as means ± SEM (\**P* < 0.05, \*\**P* < 0.01 vs. NFD + Sc Peptide; #*P* < 0.05, ##*P* < 0.01 vs. HFD + Sc Peptide; one-way ANOVA).

Changes in mRNA expression of the corresponding downstream lipogenic enzymes, regulated by the above-described transcription factors, were investigated following HFD and NFD treatments. No changes in hepatic expression of transcripts for fatty acid synthase (*Fas*), ATP citrate lyase (*Atpcl*), or acetyl-CoA carboxylase (*Acc*) enzymes were detected (Figure 6A–C). However, hepatic *Gpat3* (2.9 ± 0.5) was increased after a 6-week HFD compared with NFD feeding (0.9 ± 0.1, *P* = 0.0001) (Figure 6D). In contrast, hepatic *Gpat3* mRNA expression was significantly decreased in HFD-fed mice administered PRO20 (1.84 ± 0.18, *P* = 0.0183) compared with their control HFD counterparts. Western blotting assays confirmed modulation of GPAT3 protein by PRR antagonism, revealing that hepatic GPAT3 protein increased in HFD-fed mice (1.53 ± 0.03) mice compared with NFD controls (1.00 ± 0.18, *P* = 0.0242) (Figure 6E) and that this increase was absent in HFD-fed mice treated with PRO20 (0.93 ± 0.19, *P* = 0.0127) (Figure 6F). These data suggest that PRR antagonism regulates lipogenesis mediated by PPARγ and GPAT3, as proposed in Figure 6G.

**Figure 6.**
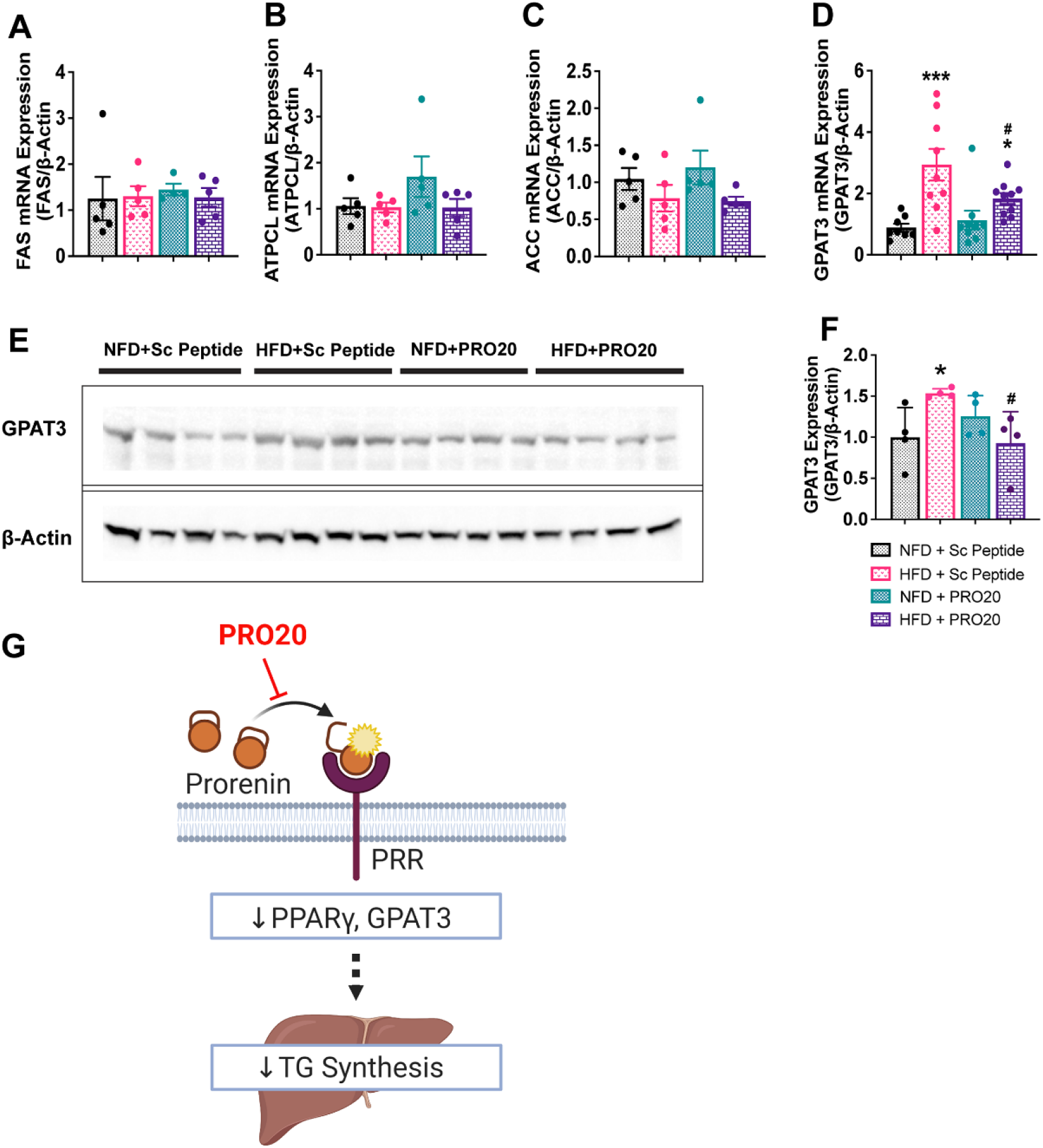
PRR antagonism prevents an increase in hepatic GPAT3 following 6 weeks of HFD. Hepatic mRNA expression of enzymatic intermediates involved in triglyceride synthesis, including FAS **(A)**, ATPCL **(B)**, ACC **(C)**, and GPAT3 **(D)**. **(E)** Western blot assessment of hepatic GPAT3 protein abundance. **(F)** Quantification of hepatic GPAT3 protein levels following 6 weeks of HFD or NFD and administration of either PRO20 or Sc Peptide. **(G)** Proposed mechanism for the reduction of liver steatosis under HFD conditions as a result of treatment with the PRR antagonist PRO20. Inhibition of PRR activation with PRO20 decreases levels of the lipogenesis transcriptional regulator PPARγ and the triglyceride synthesis enzyme GPAT3, leading to a corresponding reduction in liver triglyceride synthesis. Data are presented as means ± SEM (\**P* < 0.05, \*\*\**P* < 0.001 vs. NFD + Sc Peptide; #*P* < 0.05 vs. HFD + Sc Peptide; one-way ANOVA).

### PRR Antagonism Has no Effect on Body Weight or Glucose Homeostasis

To investigate whether PRR antagonism impacts obesity or glucose homeostasis following HFD, we monitored body weights of all mice on a weekly basis (Figure 7A). PRO20 treatment did not affect body weight in NFD-fed mice compared with scrambled peptide-treated, NFD-fed mice. After a 6-week HFD, both scrambled peptide- and PRO20-treated mice exhibited a significant increase in body weight compared with their NFD counterparts. However, no difference in body weight was observed between PRO20- and scrambled peptide-treated mice fed a 6-week HFD (Figure 7A), indicating that PRR antagonism had no effect on body weight. At baseline, there was no difference in fasting blood glucose (FBG) or glucose handling among groups, as demonstrated by glucose tolerance tests (GTTs) (Figure 7B and 7C). In contrast, at the end of the diet regimen, FBG was significantly elevated in both scrambled peptide-treated (131.4 ± 8.8 mg/dL, *P* = 0.0021) and PRO20-treated (134.9 ± 6.9 mg/dL, *P* = 0.0007), HFD-fed mice compared with their corresponding NFD-fed controls (Figure 7D). NFD-fed mice displayed no change in FBG over the duration of treatment. A GTT performed at the end of the 6-week dietary regimen (NFD or HFD) to assess the state of glucose handling showed that the HFD significantly impaired glucose tolerance (15,169 ± 786 RU, *P* = 0.0355), as reflected in an AUC analysis of blood glucose values (Figure 7F). We found no difference in glucose tolerance between HFD-fed mice treated with a scrambled peptide and those treated with PRO20 after 6 weeks. These data suggest that 6 weeks of HFD impairs glucose metabolism and promotes body weight gain, and that PRR antagonism has a minimal impact on these effects.

**Figure 7.**
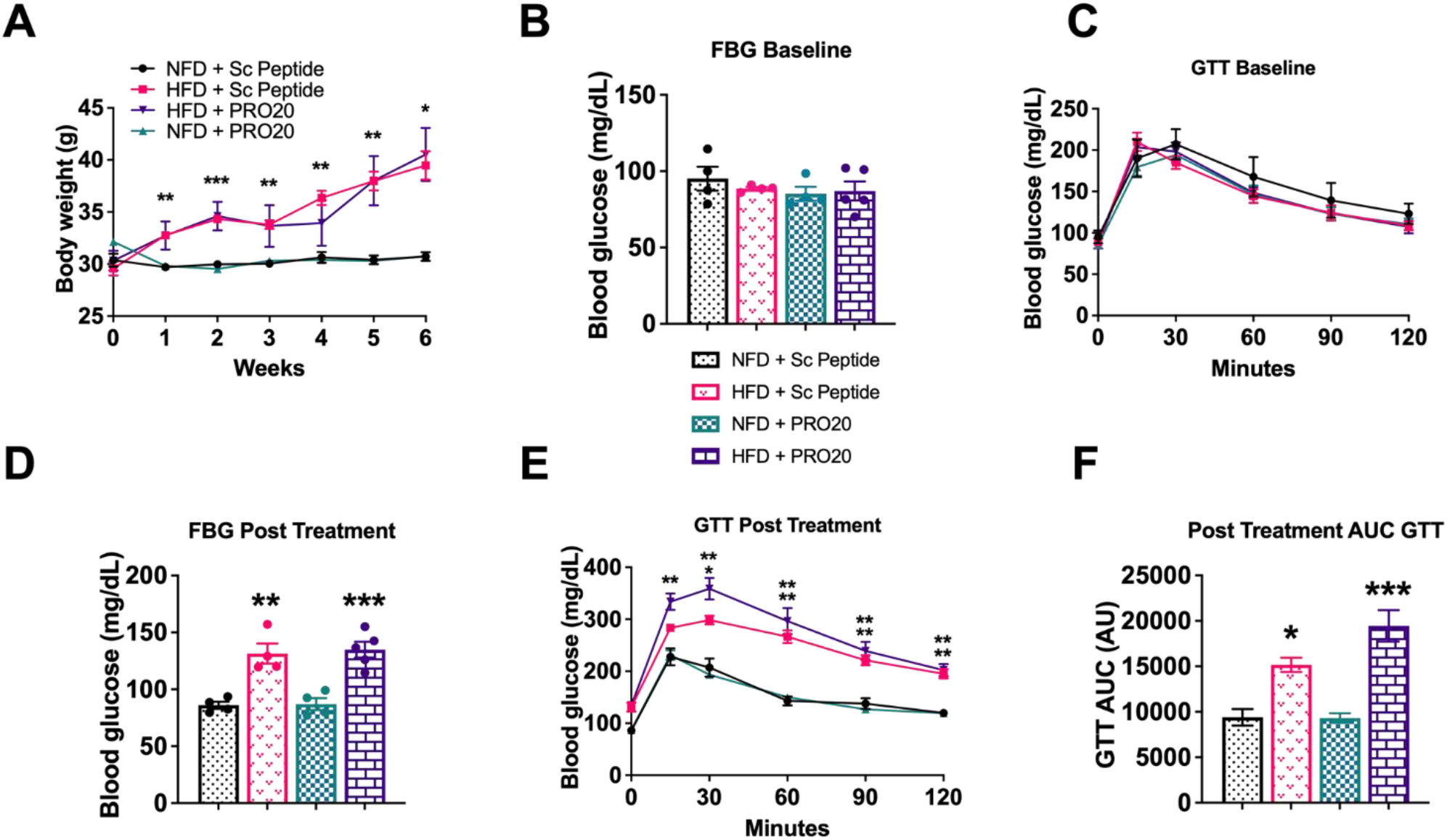
PRO20 treatment has no effect on body weight or glucose handling after 6 weeks of HFD. **(A)** Weekly body weight measurements in mice over the course of the study. **(B, C)** Baseline fasting blood glucose (FBG) levels (B) and glucose tolerance test (GTT) measurements (C) before diet modification. **(D, E)** Post-treatment measurement of FBG levels (D) and GTT measurements (E). **(F)** AUC analysis of post-treatment GTT results. Data are presented as means ± SEM (\**P* < 0.05, \*\**P* < 0.01, \*\*\**P* < 0.001 vs. NFD + Sc Peptide; one-way ANOVA).

## 4. Discussion

NAFLD is a condition in which optimal lipid-processing capabilities of the liver are altered, promoting the accumulation of fat within hepatocytes and, in later stages, the development of inflammation, culminating in necrosis and fibrosis. Despite continued increases in the prevalence of NAFLD, lifestyle modifications, though often ineffective, remain the only treatment option available and there are currently no effective pharmacological interventions for the treatment of late stages of NAFLD, including NASH with fibrosis [29]. Because of its extensive involvement in cellular metabolism in the liver, the RAS has been implicated in the pathogenesis of NAFLD, and research has demonstrated a role for RAS intermediates in hepatic glucose metabolism, lipid processing, and insulin sensitivity [30–32]. The PRR in particular has been shown to be involved in hepatic cholesterol clearance; however, the mechanisms through which the PRR regulates other aspects of lipid metabolism in the context of fatty liver and NASH development remain elusive.

In this study, we demonstrate that PRR antagonism *in vivo* ameliorates the development of HFD-induced fatty liver disease in mice. Among our key findings was the demonstration that PRO20 reduces HFD-induced lipid deposition in the liver as well as liver triglyceride content without significantly impacting body weight or glucose metabolism. We also show that losartan, an AT1R blocker, does not attenuate development of hepatic steatosis in HFD-fed mice compared with corresponding saline-treated HFD-fed controls. A further mechanistic investigation into whether modulation of the *de novo* lipogenesis pathway underlies the protective effect of PRR antagonism against lipid accumulation revealed that PRO20 treatment reduced mRNA expression of the transcription factor PPARγ and its corresponding downstream pathway enzyme GPAT3 in HFD-fed mice. Lastly, a role for the PRR in the development of fibrosis was demonstrated in MCD diet-fed mice administered PRO20, which caused a reduction in hepatic collagen deposition and serum ALT levels.

Because infusion of losartan did not reduce hepatic lipid deposition, the effect of the PRR antagonist, PRO20, in ameliorating the development of fatty liver can be explained by modulation of RAS-independent signaling pathways in hepatic lipid metabolism. Previous research has shown that the PRR contributes to the regulation of cholesterol metabolism, and that silencing of the hepatic PRR decreases low-density lipoprotein receptor (LDLR) and SORT1 protein levels in a RAS-independent manner, thereby reducing LDL clearance in the liver [9, 15]. It has also been established that hepatic PRR inhibition reduces triglyceride content, a finding also demonstrated in the current study. To narrow the mechanistic possibilities for the action of PRR inhibition on steatosis development, we performed the losartan study using diet treatment identical to that employed for the PRO20 treatment protocol. Losartan is an AT1R antagonist that competitively blocks the action of Ang II at AT1Rs at the dose employed in this study, leading to a decrease in blood pressure [33–35]. Administration of losartan has been shown to attenuate hepatic steatosis in some animal models; however, in other cases, no effect of this intervention on steatosis development, weight gain, or glucose handling was reported [36, 37]. Nevertheless, compared with the results of PRO20 intervention, losartan treatment was unable to normalize fat accumulation in the liver following 6 weeks of HFD. The lack of difference in body weight gain and glucose handling between HFD-fed mice treated with scrambled peptide versus PRO20 suggests that the PRR may target pathways separate from carbohydrate metabolism. Glucose metabolism and insulin sensitivity are key factors in the pathogenesis of NAFLD, and insulin signaling is involved in regulating fatty acid metabolism through upregulation of genes involved in *de novo* lipogenesis and downregulation of lipid-degradation pathways [39]. It is notable that obesity coupled with insulin resistance and type 2 diabetes are common comorbidities that exacerbate the progression of NAFLD in a clinical setting. In the insulin-resistant state, hepatic glucose production and fatty acid uptake are elevated, leading to increased substrates for triglyceride synthesis [40]. The fact that PRR antagonism was unable to alter body weight, fasting blood glucose or glucose handling in HFD-fed mice may suggest that our intervention only targets enzymatic pathways directly involved in lipid synthesis while leaving other clinical aspects of NAFLD pathology unchanged.

In HFD-induced non-alcoholic fatty liver development, PRR antagonism was shown to act on the triglyceride synthesis pathway by regulating PPARγ and modifying the activity of the downstream target, GPAT3. PPARγ is a transcription factor found primarily in adipose and liver tissue that modulates a multitude of target genes involved in processes such as fat storage and import, insulin sensitivity, and inflammatory response [41]. In hepatocytes, PPARγ plays a critical role in lipid homeostasis by upregulating genes involved in *de novo* lipogenesis, fat uptake, and formation of lipid droplets [41–43]. A significant increase in hepatic PPARγ expression is also a common phenotype in NAFLD models and is associated with increased steatosis [44]. Interestingly, a recent study reported that the PRR gene is a target of PPARγ *in vitro*, providing a basis for functional interactions between the PRR and PPARγ [9]. Another confirmed target of activated PPARγ is GPAT3, one of four GPAT isoforms that catalyze the conversion of glycerol-3-phosphate and long-chain acyl-CoA to lysophosphatidic acid—the pivotal rate-limiting step in the *de novo* synthesis of triglycerides in mammals [45]. GPAT3 activation in response to PPARγ has been demonstrated extensively in white adipose tissue using PPARγ agonists [46]. Previous studies have shown that downregulation of GPAT3 and other triglyceride synthetic enzymes improves HFD-induced hepatic lipid accumulation [47], underscoring the potential contribution of hepatic GPAT3 function to NAFLD. While the origin of excessive fat accumulation in the form of triglycerides in the liver varies, it has been estimated that nearly 30% of the hepatic triglycerides that accumulate during NAFLD development are sourced from *de novo* lipogenesis in the liver [2]. Decreasing the extent of *de novo* lipogenesis in the liver has considerable implications for the development of therapeutic options for NAFLD.

Finally, we investigated a potential role of the PRR in the development of NASH, a more severe stage of NAFLD, finding that PRR antagonism reduced fibrosis development in the MCD diet-induced NASH mouse model. Serum ALT levels were also reduced in MCD-fed mice treated with PRO20, indicating that PRR antagonism can decrease the severity of NASH-derived liver damage to a certain extent. ALT is commonly released from hepatocytes in response to lipid infiltrates, and subsequent cellular dysfunction and circulating ALT levels are considered indicative of liver damage [48]. However, the exact mechanisms through which the PRR acts on fibrogenic pathways remain unclear.

The approach utilized in this current study has its limitations. All experiments were performed using only adult male mice; thus, our findings are most relevant to the pathology of fatty liver in males. A 6-week HFD regimen was found to be sufficient to produce the fatty liver phenotype in male C57BL/6J mice, with 4 weeks of PRO20 intervention being able to rescue this phenotype. However, whether PRO20 would produce similar effects in other steatosis models, including high-fructose–induced steatosis, remains to be examined in future studies. In addition, the mechanism by which PRR antagonism affects liver lipid regulation was investigated only in the context of changes in the hepatic lipogenesis pathway. Other pathways, including fatty acid β-oxidation and lipid transport (through VLDL secretion) [49] could also contribute to PRR antagonism-dependent alterations in the lipid metabolism profile of the liver. Moreover, because the PRR antagonist PRO20 was administered globally via an osmotic minipump over a set period of time, its effect on hepatic lipid metabolism could also be linked to cross-talk among other organs, particularly white adipose tissue. Additional experiments to validate the action of the PRR in the liver and confirm local tissue-specific effects may require the use of liver-specific targeting.

Our results support the hypothesis that the PRR regulates lipid metabolism in the liver through modulation of *de novo* triglyceride synthesis and also plays a potential role in regulating hepatic fibrogenesis in NASH, strengthening the relevance of the PRR as a potential therapeutic target for the treatment of the NAFLD spectrum. Further studies will be aimed at elucidating the actions of the PRR in hepatic lipid degradation and transport, as well as in regulating NASH pathology and associated fibrogenesis.

### Sources of Funding

This work was supported, in part, by grants from the National Institutes of Health (NIH/NHLBI) (R01HL122770, R35HL155008) and NIH/NIGMS (1P20GM130459) to Y. Feng Earley, NIH/NHLBI (R35HL155008-01S1) to SG. Cooper, and by research programs of the Netherlands Organization for Scientific Research (TNO): ERP-Body Brain Interactions and PMC13 to R. Kleemann. The content of this manuscript is solely the responsibility of the authors and does not necessarily represent the official views of the granting agencies.

## Disclosures

The corresponding author, Dr. Yumei Feng Earley, is the inventor of the PRR antagonist, PRO20 (Patent #: PCT/US2019/045362).

## Acknowledgement

We would like to thank Drs. Nikki Trigt and Aswin Menke from the Netherlands Organization for Scientific Research for their technical assistance. Research reported in this publication utilized the Transgenic Animal Genotyping and Phenotyping Core and the High Spatial and Temporal Imaging Core facilities, supported by the National Institute of Neural Medical Sciences of the National Institutes of Health (NHI/NIGMS 1P20GM130459).

## Notes

### Competing Interest Statement

The authors have declared no competing interest.

